# Focused ultrasound to the bilateral thalamus causally modulates human cognitive attention in a frequency- and intensity-dependent manner

**DOI:** 10.1101/2025.05.27.656481

**Authors:** Kevin A. Caulfield, Christopher T. Sege, Jacob Weaver, Keith R. Murphy, Tony Passaro, Kristy Meads, Ehsan Dadgar-Kiani, Michael U. Antonucci, Cameron Good, Lisa M. McTeague

## Abstract

The centromedian nucleus of the thalamus (CMT) is a core arousal center with potential for enhancing cognitive attention. While emergent focused ultrasound can reach this area, the stimulation paradigms which may enhance function in humans remain unknown. Here we performed bilateral stimulation of the CMT using a novel ultrasound neuromodulation wearable device using 3 distinct pulse frequencies. We found that a brief 25Hz stimulation enhanced reaction time in a in a directed attention Oddball task for at least 40 minutes. Pre-post task electrocortical EEG revealed a correlation between reaction time and the prevalence of arousal related alpha and theta cortical activity. Examination of dose dependence revealed bidirectional modulation of arousal state, with lower intensities preferentially enhancing cognitive function. Pre- and post-MRI scans showed no neuroradiological changes due to stimulation. Collectively, these findings demonstrate that targeted LIFU stimulation of the CMT can safely enhance cognitive function in healthy individuals with carefully designed parameters.

## Background

The centromedian nucleus of the thalamus (CMT) plays a key role in regulating sleep-wake function^1^ and is a central hub in thalamocortical circuitry that facilitates cortical-subcortical communication^2^. Much of our understanding of CMT function has come from animal research^2,3^ or correlational neuroimaging studies in humans^4^, as its deep anatomical location has historically made it a poor target for noninvasive brain stimulation^5,6^ and precluded causal investigation of CMT function. Moreover, the ability to exogenously modulate alertness in adults could facilitate developing treatments for individuals suffering from sleep-wake disorders.^7,8^ For instance, an estimated 20% of the industrialized world works nontraditional hours, a schedule which is associated with numerous health issues such as neurocognitive impairment and increased prevalence of cancer. Shift Work Sleep Disorder^9^ affects up to 40% of individuals working this nonconventional schedule and can cause issues with falling asleep or feeling sleepy at unwanted times. Sleep disorders have also been identified in many neurological and psychiatric conditions, including depression^10,11^, anxiety^12^, and Alzheimer’s disease^13,14^, and cost an estimated $94.9 billion in the lost productivity in the US alone^8^. Thus, there is a dire need to develop treatments promoting alertness and healthy sleep-wake cycles, particularly as existing treatments such as sleep medications can cause cognitive side effects and long term dependency^15^.

We sought to test whether an emerging form of noninvasive brain stimulation – low intensity focused ultrasound (LIFU)^16,17^ – could be used to causally modulate the CMT in healthy awake adult participants. Unlike other noninvasive approaches, LIFU stimulation intensity does not decay with distance from the scalp, and it can focally target deep brain nuclei^16,17^. Animal research has shown that CMT stimulation can alter the sleep-wake behaviors in mice and rats with specific parameters^3,18^ but there have not yet been investigations in human participants that use LIFU to alter alertness in humans. Moreover, human studies have not explored critical parameters such as pulse repetition frequency^17^ for CMT to determine how this area can be optimally engaged with LIFU.

Here we conducted a within-subject, double-blind, counterbalanced, and sham-controlled study investigating arousal effects of noninvasive LIFU directed to the bilateral CMT using a newly developed ultrasound wearable. We report converging behavioral and EEG evidence of causal frequency- and intensity-dependent 25Hz LIFU-mediated changes in enhanced alertness. We also report foundational safety data demonstrating no neuroradiological changes from pre- to post-LIFU via an MRI scan battery. Together, these data suggest that 25Hz LIFU may be a safe and useful method of enhancing alertness in a non-pharmacological manner, and we present guiding parameter-related information for how LIFU could be further developed to treat sleep-wake disorders such as Shift Work Sleep Disorder.

## Results

Human participants performed a visual cognitive attentional Oddball task^19^ before and after each of 4 different LIFU parameter sets on different days in a counterbalanced, within-subject, double-blind, and sham-controlled protocol (**Figure 1A**). We tested the hypothesis that specific ultrasonic pulse repetition frequencies (PRFs) and intensities (ISPPA on target) could bidirectionally modulate the bilateral CMT using behavioral (reaction time; RT) and electrocortical (EEG) assays (**Figure 1B**). A novel wearable bilateral 128-element ultrasonic transducer (Attune Neurosciences, Inc.; **Figure 1C**) targeted the bilateral CMT with high precision and accuracy via k-wave modeling^20^ on pseudo-CT PETRA scans and 50 seconds of stimulation was delivered in each session. Sham stimulation was delivered via unfocused diffuse ultrasonic beams at the same scalp-level intensity but without focusing stimulation to the bilateral CMT (**Figure 1D**). For details see **Methods**.

**Figure 1:**
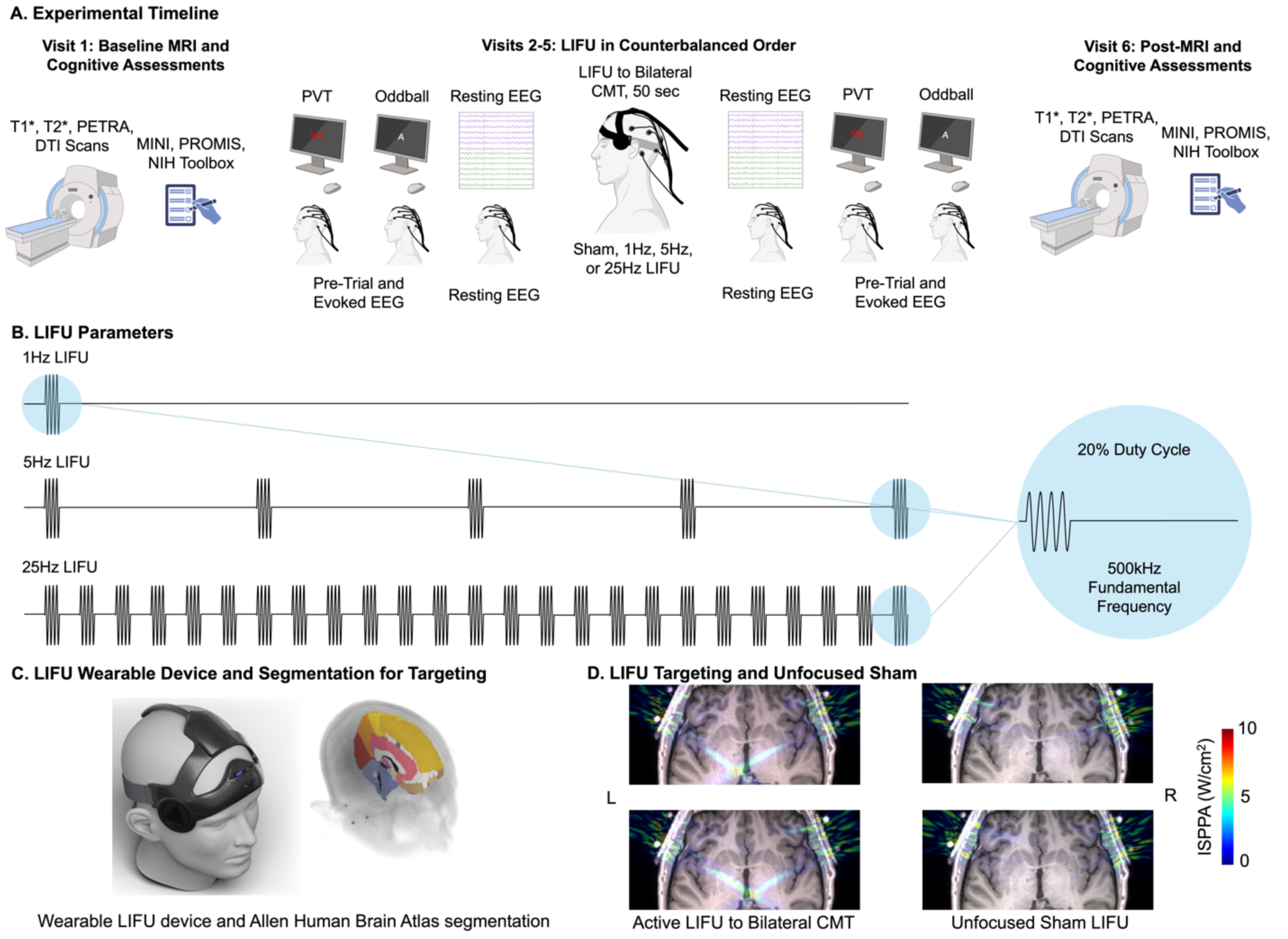
**Experimental Overview and LIFU Parameters and Simulations**. **1A.** Experimental Timeline: This study entailed 6 visits. In Visits 1 and 6, participants underwent MRI scanning and cognitive assessments. Visits 2-5 delivered LIFU at 4 pulse repetition frequencies (1Hz, 5Hz, 25Hz, unfocused diffuse sham) in a fully counterbalanced order. **1B.** LIFU Parameters showing the 3 active stimulation frequencies, duty cycle, and fundamental frequency. **1C.** LIFU Simulations showing the wearable Attune ATTN201 stimulation device and segmentation pipeline. **1D.** LIFU Targeting and Unfocused Sham for a representative participant.

### 25Hz LIFU Significantly Speeds Attentional Reaction Time

We first tested whether there is a LIFU-mediated change in attention by examining behavioral RT change on the Oddball task from pre- to post-LIFU at each PRF, focusing on trials in which participants responded accurately. We found frequency-dependent effects of LIFU such that 25Hz PRF elicited significantly faster reaction times, t(15) = 2.69, p = 0.017, d = 0.67 with an average reduction of 13.0 ± 4.9ms (**Table 1**; **Figure 2**). In contrast, there were no significant reductions in reaction times for the 1Hz, 5Hz, and unfocused diffuse sham conditions.

**Figure 2:**
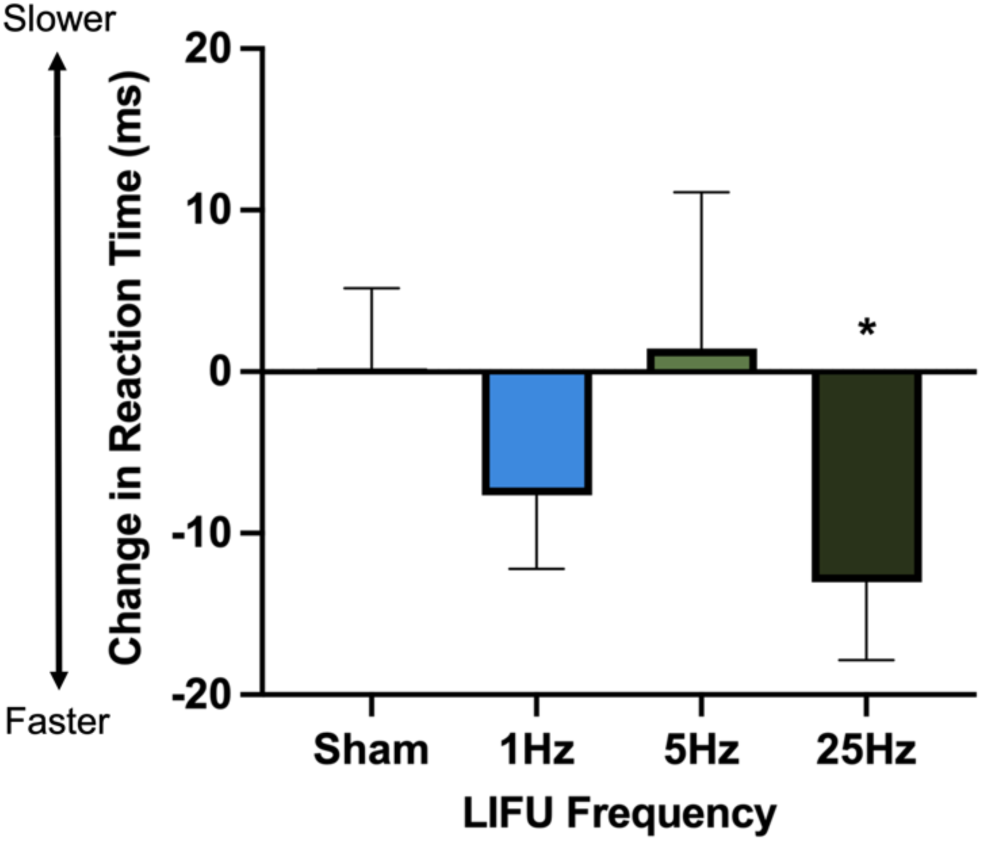
O**d**dball **Reaction Time Changes from Pre- to Post-LIFU.** We found a significant reduction in reaction time following 25Hz LIFU directed to the CMT (*p < 0.05). Change in RT is represented in ms with SEM error bars.

**Table 1:**
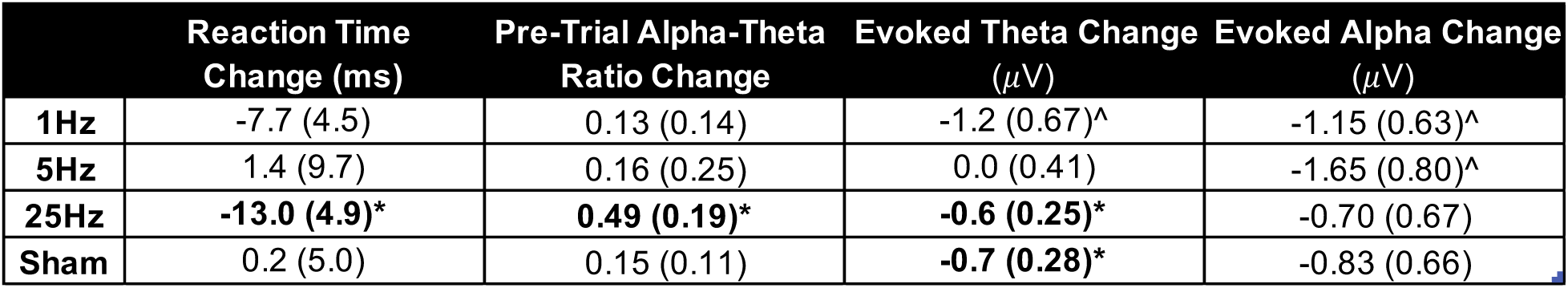
Change in Oddball Attention Assays. Data are changes from post-LIFU minus pre-LIFU; negative values indicate that the post-LIFU number was lower (i.e., faster reaction time or lower EEG amplitude or ratio). SEM is shown in parentheses. Bolded cells indicate significance; *p < 0.05, ^p < 0.10.

| | Reaction Time Change (ms) | Pre-Trial Alpha-Theta Ratio Change | Evoked Theta Change ($\mu\text{V}$) | Evoked Alpha Change ($\mu\text{V}$) |
| --- | --- | --- | --- | --- |
| **1Hz** | -7.7 (4.5) | 0.13 (0.14) | -1.2 (0.67)<sup>^</sup> | -1.15 (0.63)<sup>^</sup> |
| **5Hz** | 1.4 (9.7) | 0.16 (0.25) | 0.0 (0.41) | -1.65 (0.80)<sup>^</sup> |
| **25Hz** | **-13.0 (4.9)\*** | **0.49 (0.19)\*** | **-0.6 (0.25)\*** | -0.70 (0.67) |
| **Sham** | 0.2 (5.0) | 0.15 (0.11) | **-0.7 (0.28)\*** | -0.83 (0.66) |

Notably, the 25Hz LIFU-mediated improvements were consistent with improvements in 14 of 16 participants.

### 25Hz LIFU Significantly Alters Pre-Trial EEG Metrics of Attention

We additionally tested electrocortical neural effects during the Oddball task by conducting time-frequency decomposition of EEG data, which allowed us to test for effects in both pre-trial activity (indicative of general neural processing) and post-trial onset-evoked neural response (indicative of discrete neural processing of specific task events). Detailed analysis procedures are reported in **Methods**. Drawing on prior literature reporting that theta band (4- 8Hz) decreases^23^ and alpha-band (8-12Hz) increases^24^ in non-task EEG are associated with heightened alertness, and that a ratio of these oscillatory signatures can capture cognition-related brain activity^25,26^, we focused on a pre-trial alpha-theta ratio at pre- and post-LIFU timepoints as a primary index of electrocortical changes due to LIFU.

In the 500ms pre-trial period, we found a significant increase in frontal alpha-theta ratio from pre- to post-25Hz LIFU that were in line with increases in alertness, t(14) = 2.54, p = 0.024, d = 0.66, with an average increase in the ratio of 0.492 ± 0.193 and an increase in alpha-theta ratio present in 12 of 15 participants (**Figure 3A-D** and **Table 1**). In contrast, there were no significant LIFU-induced changes in alpha-theta ratio after 1Hz, t(14) = 0.868, p = 0.40, d = 0.22, 5Hz, t(14) = 0.64, p = 0.53, d = 0.17, or sham, t(14) = 1.33, p = 0.21, d = 0.34, stimulation.

**Figure 3:**
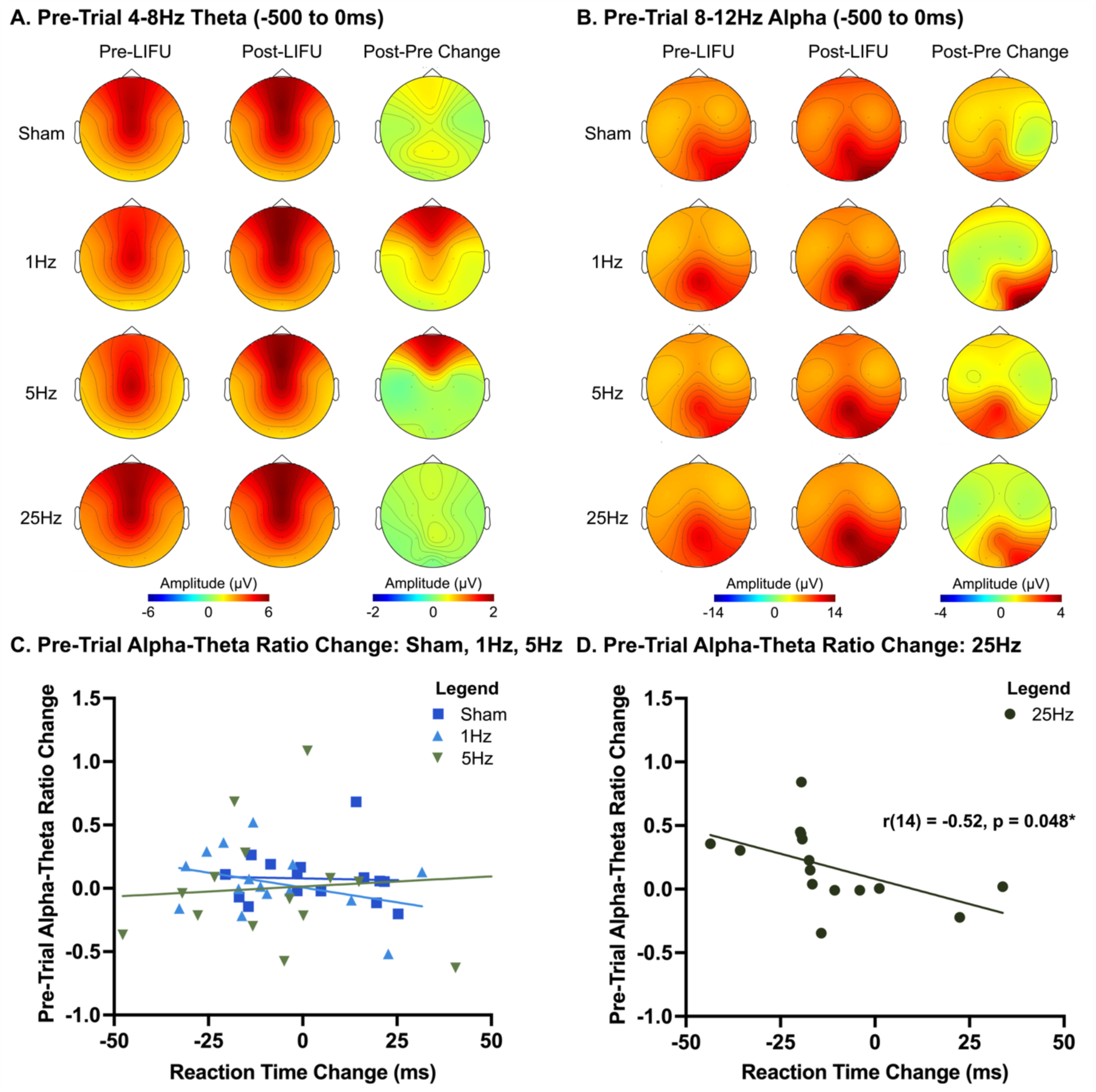
O**d**dball **Pre-Trial (-500 to 0ms) Alpha and Theta Activity as a Function of RT. 3A-B.** Pre-LIFU, Post-LIFU, and Post-Pre Change in Theta (A) and Alpha (B) EEG Signal. **3C.** Pre-Trial Alpha-Theta Ratio Change for Sham, 1Hz, and 5Hz Conditions. **3D.** Pre-Trial Alpha-Theta Ratio Change for 25Hz. We found a significant increase in alpha-theta ratio from pre- to post-LIFU in the 25Hz LIFU condition but not the 1Hz, 5Hz, or unfocused diffuse sham conditions. This 25Hz LIFU-mediated increase in alpha-theta ratio correlated with changes in RT on Oddball target trials directly linking the brain and behavioral changes induced by 25Hz LIFU targeting the bilateral CMT.

Further substantiating our hypothesis that LIFU can modulate human attention, we found a significant Pearson’s correlation between change in RT and change in pre-trial alpha-theta ratio in the 25Hz LIFU condition, r(14) = -0.52, p = 0.048 (**Figure 3D**). This relationship was not present for any of the other stimulation frequencies, including 1Hz (r(14) = -0.344, p = 0.21), 5Hz (r(14) = 0.136, p = 0.63), and sham conditions (r(14) = -0.042, p = 0.88). These data suggest a direct relationship between 25Hz LIFU-induced changes in oscillatory rhythms and increased alertness – i.e., behavioral gains indicative of enhanced attention.

### 25Hz LIFU Significantly Alters Evoked EEG Metrics of Attention

In addition to pre-trial electrocortical activity, we examined how EEG activity changes after trial onset in the Oddball task changed from pre- to post-LIFU (**Figure 4A-D**). We again focused on evoked changes in theta (4-8Hz) and alpha (8-12Hz) EEG bands that occurred in sensor/ time windows consistent with prior literature, i.e., 100-550ms for a frontal theta “burst” and 150-600ms for a parietal alpha “block.” In contrast to pre-trial activity, evoked EEG *increases* in theta and active *blocking* of alpha activity are associated with increases in attention.

**Figure 4:**
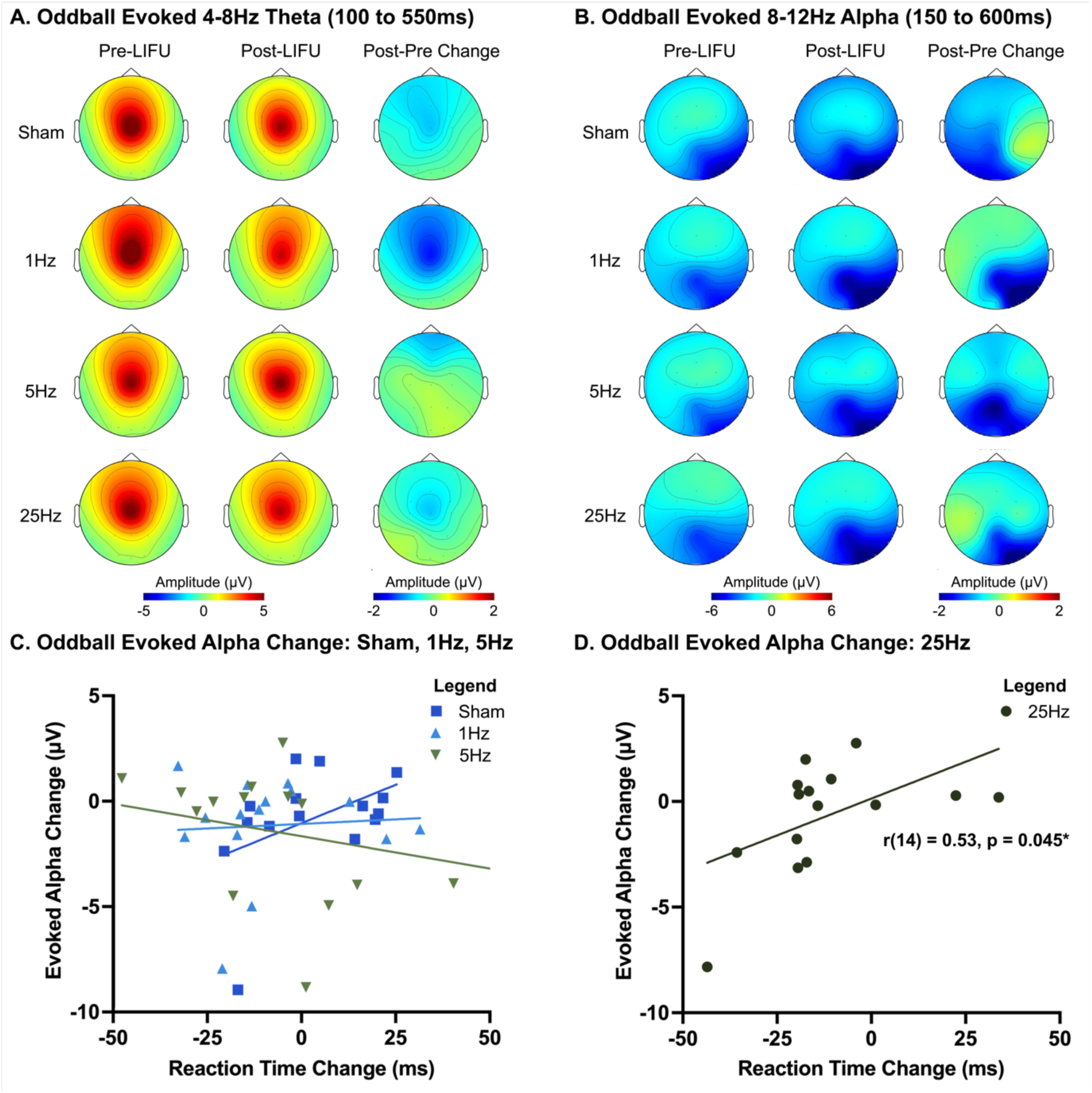
O**d**dball **Evoked Alpha and Theta Activity and Correlations with RT. 4A-B.** Pre-LIFU, Post- LIFU, and Post-Pre Change in Evoked Theta (A) and Evoked Alpha (B) EEG Signal. **4C.** Evoked Alpha Change Correlations for Sham, 1Hz, and 5Hz Conditions. **4D.** Evoked Alpha Change Correlation for 25Hz LIFU. We report a significant correlation between evoked alpha change (μV) and reaction time change (ms) in the 25Hz LIFU but not other conditions, such that participants with more substantial reductions in evoked alpha tended to also have a greater reduction in reaction time from pre- to post-LIFU.

Within this context, we found decreases in the evoked theta response from pre- to post-LIFU across 25Hz, t(14) = 2.45, p = 0.028, d = 0.63, sham, t(14) = 2.228, p = 0.043, d = 0.58, and (at a trending level) 1Hz, t(14) = 1.86, p = 0.08, conditions (**Table 1**); but, this effect was not present in the 5Hz condition, t(14) = 0.06, p = 0.955, d = 0.015. Regarding evoked alpha blocking, then, there were no significant changes in this response from pre- to post-LIFU for 25Hz, t(14) = 1.04, p = 0.32, d = 0.27, or sham conditions, t(14) = 1.245, p = 0.23, d = 0.32, and there were trending decreases in alpha blocking in the 1Hz, t(14) = 1.82, p = 0.09, d = 0.47 and 5Hz condition, t(14) = 2.07, p = 0.057, d = 0.53.

In correlations relating pre- to post-LIFU change in evoked alpha with RT values, we again found a significant correlation in the 25Hz LIFU condition, r(14) = 0.53, p = 0.045, indicating that participants with a more substantial increase in evoked alpha blocking after LIFU also had a greater improvement (i.e., reduction) in reaction time (**Figure 4D**). This relationship was not present for the 1Hz, r(14) = 0.065, p = 0.82, 5Hz, r(14) = 0.39, p = 0.15, or sham, r(14) = 0.444, p = 0.097, conditions. In conjunction with the 25Hz LIFU-induced changes in pre-trial alpha-theta ratio change, then, it appears that brief 25Hz LIFU substantially improves directed attention and that this was caused by changes in task-associated neural oscillatory activity presumably supported by the targeted thalamic node.

### Inverse Relationship Between ISPPA Intensity and Oddball Neurobehavioral Effects

Given significant attention and electrocortical changes elicited by active 25Hz LIFU, we examined whether the on-target ISPPA stimulation intensity (averaged across the left and right CMT) correlated with pre- to post-LIFU changes. The average on-target ISPPA values in the bilateral CMT were 4.37 ± 0.31 W/cm^2^ (range = 2.69-6.27 W/cm^2^). Lowest, median, and highest ISPPA images are displayed in **Figure 5A**.

**Figure 5:**
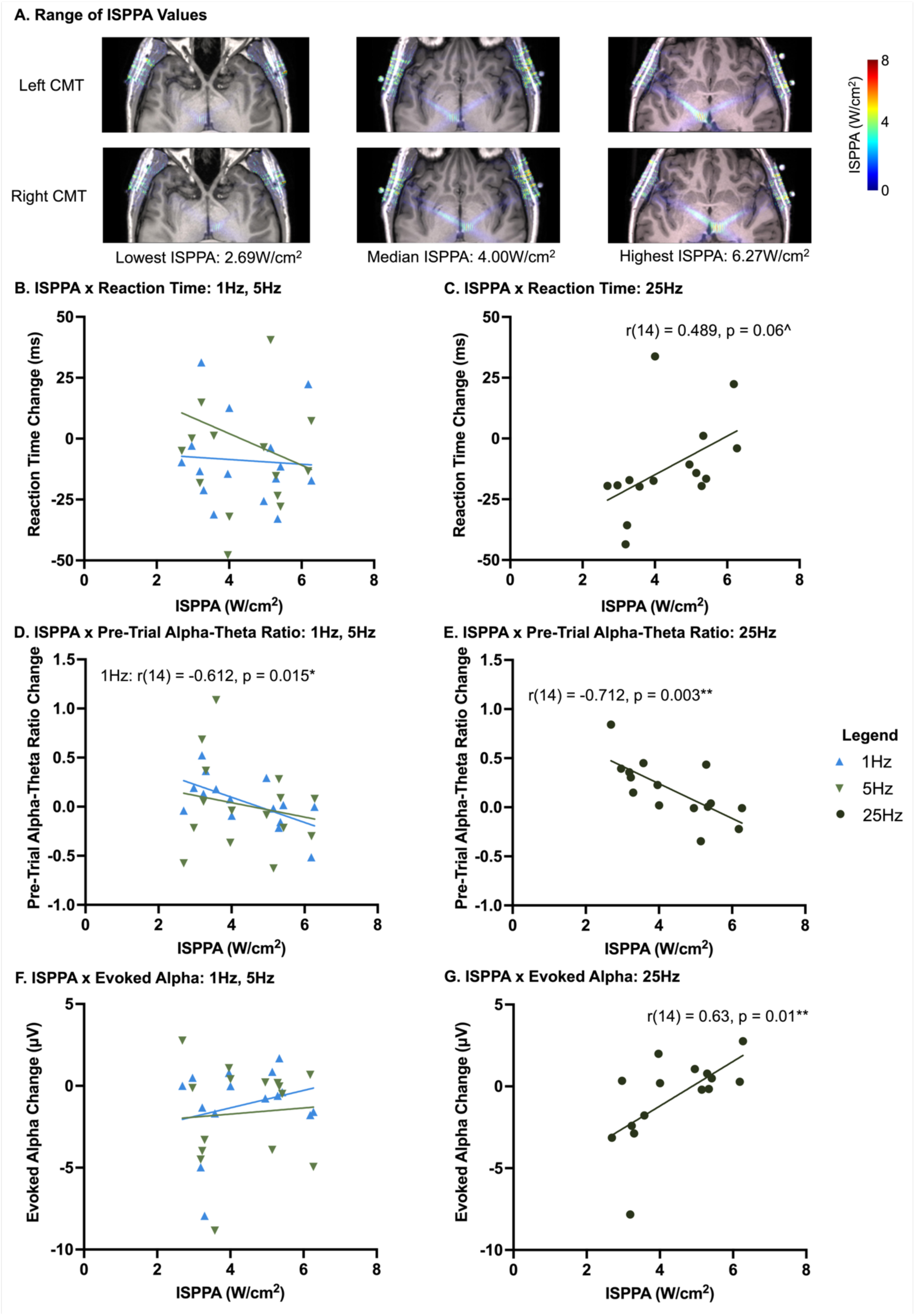
I**S**PPA **Values and Correlations with Oddball RT, Oddball EEG Pre-Trial Alpha-Theta Ratio, and Oddball EEG Evoked Alpha. 5A.** Range of ISPPA values for the Left and Right CMT. **5B-C:** ISPPA x Oddball RT. There was a trending relationship between lower ISPPA values eliciting the largest reductions in reaction time from pre- to post-25Hz LIFU (p = 0.06^). **5D-E:** ISPPA x Oddball Pre-Trial Alpha-Theta Ratio. There were significant correlations between ISPPA x pre-trial alpha-theta ratio from 25Hz (p = 0.003**) and 1Hz LIFU (p = 0.015*) but not 5Hz. 5F-G: ISPPA x Evoked Alpha. There was a significant relationship between ISPPA and evoked alpha change such that the greatest reductions in evoked alpha were associated with lower ISPPA values (p = 0.01**). This relationship was not present in the 1Hz or 5Hz LIFU conditions.

For reaction time, we found a trending correlation between ISPPA and RT to targets for 25Hz LIFU (**Figure 5C**), r(14) = 0.489, p = 0.06; such that greater improvements in RT were associated with lower ISPPA intensities. Conversely, we did not observe this relationship for 1Hz (r(14) = 0.064, p = 0.82) or 5Hz conditions, (r(14) = 0.199, p = 0.48; **Figure 5B**). We did not compute correlations for the sham condition as, by definition, ISPPA values were consistently near 0W/cm^2^.

Similarly, stronger EEG changes were associated with lower ISPPA intensities. For pre-trial alpha-theta ratio change from pre- to post-LIFU we found a significant inverse relationship with ISPPA for the 25Hz condition, r(14) = -0.712, p = 0.003 (**Figure 5E**). In addition, a correlation in the same direction between ISPPA values and pre-trial alpha-theta ratio change was significant for the 1Hz condition, r(14) = -0.612, p = 0.015; however, this was not the case for 5Hz, r(14) = -0.194, p = 0.51 (**Figure 5D**). In the same vein, for pre- to post-LIFU change in evoked (i.e., post-trial onset) alpha blocking, we again found a significant correlation with ISPPA for 25Hz LIFU, r(14) = 0.63, p = 0.01 (**Figure 5G**), such that larger increases in evoked alpha blocking were associated with lower ISPPAs. Both the 1Hz condition, r(14) = 0.26, p = 0.35, and the 5Hz LIFU condition, r(14) = 0.073, p = 0.80, did not have significant relationships between ISPPA and evoked alpha signal (**Figure 5F**). Collectively, this converging evidence indicates that larger changes in alertness indicators were associated with *lower* intensities of 25Hz LIFU, across RT and EEG indices.

### Safety Profile: MRI Scans and Tolerability

To evaluate potential structural changes or observable complication associated with LIFU, a board-certified neuroradiologist conducted a blinded review of MRI scans obtained both before and after 4 sessions of LIFU. Given the novel nature of this technology, the imaging protocol included both standard anatomical sequences (volumetric T1 MPRage) and additional sequences to assess for potential complications. Specifically, the scan battery consisted of: diffusion-weighted imaging to exclude infarct/ischemic injury; T2-susceptibility imaging to assess for parenchymal hemorrhage or blood breakdown products; 3D T2-FLAIR to identify edema, new signal abnormality, and hemorrhage in the extra-axial or sulcal spaces; and volumetric T2- weighted imaging to evaluate morphological change in ventricular or CSF space anatomy. Particular attention was given to the region of ultrasound stimulation consisted of an evaluation of the underlying scalp, calvarium, extra-axial space and brain parenchyma. Pre- and post-stimulation MR examinations were reviewed in a randomized side-by-side fashion. Thus, the radiologist was unaware of the timing of the MRI relative to ultrasound stimulation. The presence or absence of difference between the paired images was recorded.

No structural abnormalities or imaging changes were observed. Specifically, there was no parenchymal signal abnormality suggestive of infarct, injury, inflammation, or hemorrhage. No extra-axial blood was identified. No change in the calvarium or scalp was observed in the region of device placement (or elsewhere).

This safety profile dovetails with our tolerability findings reported in detail in **Supplemental Section 1**, in which we found that headache, pain, skin/scalp irritation, facial twitching, fatigue, and fearfulness/anxiety were very low with all values at an average of 1.18 out of 10 or lower on a scale of 0 (low) to 10 (high) and not persisting beyond the stimulation period.

## Discussion

To our knowledge, this is the first human study reporting significant increases in alertness via LIFU stimulation to the bilateral CMT. In converging lines of evidence from behavioral (reaction time) changes and electrocortical recordings (alpha and theta EEG), we found that 50 seconds of 25Hz LIFU stimulation significantly enhanced performance on a directed attention task up to at least 40 minutes post-stimulation. These improvements in reaction time significantly correlated with neural changes in both task and non-task epochs, directly linking together brain and behavioral changes. However, enhanced alertness from LIFU was frequency-dependent and restricted to the 25Hz LIFU frequency, as we did not observe significant changes across behavioral and electrocortical assays in the 1Hz, 5Hz, or unfocused diffuse sham conditions. Moreover, we report that ISPPA intensity at the bilateral CMT inversely correlated with 25Hz-mediated changes in alertness such that the largest gains were in participants receiving the lowest intensity values. These data have several implications: 1) The bilateral CMT plays a causal role in human alertness; 2) 25Hz LIFU to the bilateral CMT can enhance alertness; 3) Thalamic ultrasound can elicit electrocortical changes in neural activity detected by EEG; 4) 3x sessions of active LIFU is safe and well-tolerated, as determined by neuroradiological reads and questionnaires. Taken as a whole, 25Hz LIFU to the bilateral CMT causally enhances alertness in a frequency- and intensity-dependent manner.

Contextualizing these findings within the literature, this study demonstrates that LIFU to the CMT can modulate cortical activity in a manner consistent with heightened alertness. Prior studies suggest that resting frontal theta oscillations are indicative of an increase in drowsiness and sleep pressure, as evidenced by an increase in theta activity during sleep deprivation in animals^21^ and humans^22^. Furthermore, some researchers have hypothesized that increased tonic (i.e., not task event-evoked) theta activity represents a type of localized sleep during wakefulness^23^. Thus, our finding of a 25Hz LIFU-induced reduction in pre-trial theta oscillations is in line with increased alertness.

Furthermore, these data highlight the potential therapeutic opportunity of applying LIFU to the CMT to increase alertness in individuals with diagnoses such as Shift Work Sleep Disorder. Shift Work Sleep Disorder is characterized by excessive daytime sleepiness and difficulty staying awake at work, and it affects millions of people worldwide. As we found converging evidence that 25Hz LIFU can enhance alertness, a next step may be to test the effects of stimulation in individuals suffering from Shift Work Sleep Disorder and evaluating whether this may enhance their sleep-wake cycles and job performance.

These data additionally speak to the importance of considering LIFU stimulation frequency and intensity in determining the directionality of effects. We report that 25Hz LIFU increased alertness across several behavioral and electrocortical metrics whereas 1Hz, 5Hz, and unfocused diffuse sham stimulation did not. Of particular interest, prior reports using 5Hz “theta-burst” stimulation and motor physiology has led to widespread adoption of this stimulation frequency and reports of 5Hz LIFU having excitatory effects^31^. As we did not find evidence of 5Hz LIFU enhancing reaction time or EEG measures consistent with increased alertness, it is reasonable to consider whether the effects of LIFU on cortical tissue are the same as those experienced at deeper brain regions and if stimulation frequency differentially affects various brain regions. Similarly, as we applied the same LIFU intensity at the scalp-level, there was a distribution of the ISPPA intensity at the bilateral CMT between 2.69-6.27W/cm^2^ and we report that participants receiving lower ISPPA values tended to have greater changes in alertness metrics. While paradoxical, these data corroborate the preclinical animal literature in which researchers have reported that higher ISPPA intensities from 20Hz LIFU over the CMT did not result in higher fiber photometry signal; rather, for high frequency (20Hz) LIFU over the CMT, there was a non-linear relationship between intensity and brain signal^3^. In contrast, when these researchers tested for the same dose-response relationship between ISPPA intensity and fiber photometry signal from a lower PRF (2.5Hz) LIFU to the CMT and other brain regions (dorsomedial hypothalamus and locus coeruleus), there was in fact a linear relationship between intensity and fiber photometry signal (i.e., higher intensity = higher fiber photometry signal). In sum, these data highlight the importance of considering how LIFU stimulation frequency, intensity, and brain target interact. While more evidence must be collected, optimal LIFU parameters may be specific to individual brain regions.

There were some limitations in this study. We applied a 50s stimulation duration for each person, but many studies use longer stimulation durations that have been linked to stronger corticomotor effects. In addition, there is room for further optimization of other LIFU parameters beyond pulse repetition frequency and ISPPA on-target intensity. Due to the nature of the wearable stimulation device, we collected EEG using only 16 electrodes distributed throughout the head, whereas many studies use denser electrode arrays that are more capable of determining topography of brain signals. Future research may consider further optimizing stimulation parameters and acquiring EEG using additional electrodes.

## Conclusions

25Hz LIFU applied to bilateral CMT elicited behavioral (RT) and electrocortical (EEG) changes indicative of increased alertness, and lower ISPPA intensity elicited the largest changes from pre- to post-LIFU. Findings support the use of LIFU to modulate deep brain structures and provides guiding data on parameter selection.

## Methods

### Participant Demographics

Participants were 16 healthy adults (6 M, 10 F, mean age = 25.1, age range = 22-29) with no psychiatric diagnosis as confirmed by a structured interview (MINI),^24^ recruited via emailed broadcast message. Participants also completed NIH Patient-Reported Outcomes Measurement Information System (PROMIS) measures^19^ and scored in normal ranges on depression, fatigue, and cognitive ability scales.

### Study Design

This study was approved by the MUSC IRB, and consisted of 6 within-subject, double-blind, sham-controlled in-person visits (**Figure 1**). In Visit 1, participants underwent baseline structural MRI scanning and cognitive and neuropsychiatric testing (MINI Mental Status Exam^24^, NIH Cognition Toolbox^25^, PROMIS measures^26^). Visits 2-5 comprised of a 50s LIFU stimulation period with pre- and post-stimulation behavioral measures of PVT and Visual Oddball tasks in concert with EEG recording, as well as resting EEG (the latter data to be reported in a separate paper). The experimental manipulation between Visits 2-5 was the LIFU stimulation frequency, in which, at each session, participants received either unfocused diffuse sham, 1Hz, 5Hz, or 25Hz LIFU in a counterbalanced order. On each day, participants rated side effects at pre-LIFU, during-LIFU, and post-LIFU (i.e., following post-LIFU resting EEG, PVT, and Visual Oddball tasks). At Visit 6, participants then underwent the same MRI scanning and cognitive testing, which served as the post-LIFU safety and function check. These post-treatment assessments were not designed to capture individual LIFU frequency effects but, rather to measure any global impacts of the 4 visits.

### LIFU Stimulation Device and Parameters

LIFU was administered using the wearable ATTN201 device (Attune Neurosciences, Inc., San Francisco, CA, USA) which contains bilateral 64-element transducers (128 elements total). Stimulation was delivered at a 20% duty cycle, 60V/pulse, and a 500kHz fundamental frequency, using image-guided targeting of the left and right CMT with pseudoCT created from PETRA MRI scans and k-wave^20^ simulations. For sham stimulation, an unfocused diffuse field was administered with on-target ISPPA values approximately 80x lower than those from the focused CMT simulation.

For each condition, stimulation was delivered at a 20% duty cycle at 60V/pulse at the scalp level, corresponding to left CMT ISPPA values of 4.37 ± 0.32W/cm^2^ (range = 2.70- 6.45W/cm^2^) and right CMT ISPPA values of 4.37 ± 0.33W/cm^2^ (range = 2.68-6.27W/cm^2^) on average. For ISPPA analyses in **Figure 5**, we averaged the within-subject left and right ISPPA values as we stimulated the bilateral CMT, for an average of 4.38 ± 0.31 W/cm^2^ (range = 2.69- 6.27W/cm^2^). The average difference between left and right CMT ISPPA was of 0.002 ± 0.18W/cm^2^.

In all participants, the MI and thermal index values were below the US FDA limits of MI < 1.9 and temperature rise < 2° C in soft tissue. More specifically, the MI values were, in the left CMT: 0.526 ± 0.019 (range = 0.417-0.645) and in the right CMT: 0.525 ± 0.019 (range = 0.415- 0.648). Regarding temperature rise, the values in the left CMT were: 0.297 ± 0.019° C (range = 0.191-0.40° C) and in the right CMT were: 0.293 ± 0.016° C (range = 0.52-1.31° C).

### CMT Targeting Procedures

All MRI scans were acquired on a Siemens Magnetom Prisma 3T scanner (Siemens, Erlangen, DE) using a 32-channel head coil. At Visit 1, participants were fitted with an MRI-compatible version of the wearable 128-element ATTN201 device (**Figure 1C**). Using a custom 3D-printed tool, we measured and added foam pads between 0.16 and 1.58cm at four positions on the wearable device to ensure that it precisely fit each person’s head and would not move during the experimental visits. The MRI-compatible device does not contain internal electronics but does have four bilateral fiducial markers (8 total) at set positions that enabled co-registration of the device relative to each participant’s deep brain nuclei. Fiducial markers are placed over the transducer positions on the left and right temples and are visible in the PETRA (UTE) and T1* MPRAGE scan sequences. Using these markers and knowing their positioning relative to each of the 128 focused ultrasonic elements, a trained algorithm extracted their locations relative to the participants anatomy and calculated the firing sequence of each element to most accurately stimulate the left and right “Centromedian Nucleus of the Thalamus” location in the Allen Human Brain Atlas^27^.

For targeting, PETRA scans were normalized to the soft tissue (1) and air peak (1) values and converted to a pseudoCT. Using a previously reported approach, sound speed and density were derived from the pseudoCT using linear mapping, and a homogenous attenuation factor was assigned to the skull to account for pressure loss. We calibrated acoustic simulation peak free-field pressures to measured free-field peak coordinates across the target distribution range with an HGL-0200-calibrated hydrophone in degassed water (Onda Corporation, Sunnyvale, CA, USA). We used the k-wave toolbox implementation of a k-space pseudospectral time domain model of acoustic propagation and the Penne’s Bioheat simulation of temperature evolution to perform acoustic and thermal simulations. All protocol acoustic and thermal simulations yielded a Mechanical Index (MI) < 1.9 and temperature rise < 2° C in soft tissue in line with US FDA limits and consensus safety guidelines^17^. See **Figure 5A** for representative simulations showing the range of ISPPA values in this study.

In Visits 2-5, the 128-element ATTN201 LIFU device was positioned on each person’s head using the same approach and using the same foam pad spacers as at the MRI visit. The stimulation device was only turned on for the 50s stimulation period to minimize any potential electrical interference with the external EEG recordings. Individualized stimulation files with the precise firing pattern and timing of element firing were used to stimulate the left and right CMT given each person’s anatomy.

### Unfocused Diffuse Sham LIFU

An innovation that we highlight in this study is the use of an unfocused diffuse sham stimulation approach. The benefit of this method is that we delivered the same 60V/pulse at a 25Hz frequency for the same 50s duration such that all parameters are identical to the active 25Hz condition, thus producing the same sensations within intermediary tissue and nerve fibers. However, the unfocused diffuse sham condition does not focus ultrasound waves within the brain, but instead elements are fired in a deprogrammed sequence that achieves the same sensations at the scalp without meaningfully travelling to any specific brain region. In **Figure 1C**, we show the left and right CMT simulations compared to the unfocused diffuse sham condition in a representative participant to demonstrate the difference in stimulation intensity at the bilateral CMT between the two conditions.

### Visual Oddball Task: Directed Attention

In Visits 2-5, participants completed pre- and post-LIFU Visual Oddball tasks^19^ to measure changes from 50s of stimulation. The Oddball task is a measure of directed attention in which participants respond to letters rapidly presented on a computer screen. Prior to a block of 40 trials, participants were given a target letter for that block (A, B, C, D, or E; each letter serves as target once across 5 task blocks). Across 5 total blocks, 200 trials are presented at each timepoint. On each trial, participants indicated if the letter currently on the screen matches the target by pressing the “UP” arrow or does not match using the “DOWN” arrow on the keyboard. Each letter was presented for 200ms and followed by a 1100 to 1500ms jittered inter-stimulus interval). Of the 40 letters in each block, 8 are target letters and 32 are non-target (“distractor”) letters. In this study, we focused our analyses on the target trials as the Oddball task is designed to examine behavioral and neural responses to target stimuli with non-target “distractors” serving as a comparison condition. Distractor analyses are presented in **Supplemental Section 4**. To analyze behavior, we measured the RT to correct trial responses and ensured that overall accuracy between pre- and post-LIFU timepoints, as well as accuracy with different frequencies, were not significantly different.

For neural effects, prior studies indicate different neural activity correlates of enhanced alertness depending on whether activity is indexed in the pre-trial period (as a measure of tonic activity) or in the period after trial onset (to measure activity changes due to trial occurrence, i.e., “evoked” EEG). In a pre-trial period in which participants are not actively performing the task but within the context of the task period, prior data suggest enhancements in alertness track with increases in alpha activity^28^ and decreases in theta activity^29^. Given this reciprocal relationship, a number of studies compute a ratio of tonic alpha/ theta power as an integrated indicator of general alertness^30,31^. In contrast, for evoked EEG activity punctate *increases* in frontal theta activity^32^ and an active *decrease* of parietal alpha activity^33^ (alpha “blocking”) have then been characterized as indices of complementary but distinct task processing operations. Integrating these converse relationships (e.g., association of alertness with increased alpha in the pre-trial period and alpha *blocking* after trial onsets), a common contemporary interpretation is that trial processing efficiency/ effectiveness is most directly correlated with the degree which oscillatory processes are *synchronized* or *desynchronized* by trial occurrence. As such, with alpha signal, adaptive desynchronization could be enhanced by increasing pre-trial power, decreasing post-onset power, or both (whereas this adaptive contrast would not be enhanced by reducing pre-trial *and* post-trial alpha)^34^.

Consistent with distributions reported in prior literature and with observed distributions in the current study, we quantified frontal theta effects (pre-trial and evoked) in Fz, FC3, FC4, and Cz electrodes, and parietal (pre-trial and evoked) alpha across Cz, P3, P4, and Pz electrodes. For pre-trial activity, we focused on the ratio of pre-trial alpha/theta activity changes from pre- to post-LIFU in the frontal electrodes as an integrated measure of tonic alertness.

### Psychomotor Vigilance Task (PVT): Sustained Attention

Participants also completed the Psychomotor Vigilance Task (PVT)^35^ at pre- and post-LIFU timepoints as a measure of sustained attention. Results are reported in **Supplemental Section 2**. In the PVT, participants fixate on a black screen until a millisecond (ms) counter appears for up to 5000ms in red text on the center of the screen. Participants click the left mouse button as quickly as possible in response to seeing the counter appear on the screen.

After each response, the counter immediately stops, and RT is then shown for 1s to encourage rapid responding. After each counter, a jittered 1000-10,000ms inter-stimulus interval (black screen) precedes the next counter for ∼96 total trials on average over a total period of exactly 10 minutes for each subject. Per standard procedures^35^, aberrant responses are coded as “FALSE” (i.e., responses in the pre-trial period), “MISS” (i.e., no response during the counter), and “LAPSE” (i.e., response time later than 500ms). For neural responses to counters, we focused on the same cluster of frontal theta (Fz, FC3, FC4, and Cz) and parietal alpha (Cz, P3, P4, and Pz) electrodes as in the Oddball Task.

### EEG Acquisition and Analysis

We acquired EEG from 16 standard actiCHamp electrodes (Brain Products GmbH, Gilching, DE) placed at International 10-20 system sites according to a modified typical 16- channel layout (F3, Fz, F4, FC3, FC1, FC2, FC4, C3, Cz, C4, P3, Pz, P4, O1, Oz, and O2)^36^.

The typical 16-channel layout was slightly modified at frontal sites to accommodate placement of the ultrasound device. Electrodes were directly placed on the head and held on using conductive EEG paste (Ten20 Conductive Paste, Weaver & Co, Aurora, CO, USA) and were positioned using neuronavigation software^37^ (Brainsight, Rouge Inc., Montreal, CA) and MNI coordinates for each electrode to reliably position them on the participant’s head^38^. When the electrodes were placed on the head, they were resampled to ensure individual reproducibility between sessions to acquire the most accurate EEG signal as possible. Bilateral mastoid electrodes served as references and an electrode placed on the middle of the forehead was used as the ground electrode. In all visits, all electrodes had impedances of 10kΟ. Data were acquired at 5000Hz and then down-sampled to 250Hz.

Subsequent pre-processing followed standard steps executed with the FieldTrip Matlab toolbox^39^, and included: bandpass filtering from 1 – 40Hz; removal of eye movement artifact using Independent Components Analysis; visual removal of other (e.g., motion) artifacts, and; segmenting (2000ms pre-trial to 2000ms post-trial) and organizing trials by condition. Following processing, trials within each condition were submitted to time-frequency decomposition using multi-taper convolution prior to fast Fourier transform within moving windows (window length decreased as frequency increased)^40^ across the length of each trial. For pre-trial analyses, time-frequency data were averaged from -500 to 0ms pre-trial onset. For evoked EEG analyses, then, data were first baseline corrected (deviated from the 500ms pre-trial period) and then epoched in time windows of observed response maxima with 0ms representing trial (i.e., Oddball stimulus) onset. For frequency band analyses, we used conventional thresholds of 4- 8Hz for theta activity and 8-12Hz for alpha activity.

### Neurocognitive Assessments

We used several cognitive and global health assessments in Visits 1 and 6 to assess for gross changes following 4 sessions of LIFU. The PROMIS consists of self-rated scores that are fully corrected t-scores for an average of 50 ± 10 on measures of anxiety, depression, fatigue, sleep, and cognition. We used the NIH Cognition Toolbox to probe various cognitive functions across crystallized and fluid cognition domains, with fully corrected t-scores with an average of 50 ± 10. Finally, we administered the Mini-International Neuropsychiatric Interview (MINI) to confirm the absence of mood, anxiety, psychotic, or substance use disorders. See **Supplemental Section 3** for details.

### Statistical Analyses

Statistical measures consisted of t-tests, ANOVAs, and Pearson’s correlations. To test the pre- to post-LIFU behavioral and electrocortical effects of stimulation within each stimulation frequency, we utilized paired t-tests. To assess the side effects at three timepoints, we used repeated-measures ANOVAs with the repeated timepoints of pre-, during-, and post-LIFU. To evaluate the relationship between EEG and RT, as well as ISPPA and EEG and ISPPA and RT, we computed Pearson’s correlations. For all statistical tests, the significance level was set at two-tailed p < 0.05.

## Supporting information

Supplemental Sections 1-4

## References

1. Gent TC, Bandarabadi M, Herrera CG, Adamantidis AR. Thalamic dual control of sleep and wakefulness. Nature neuroscience. 2018;21(7):974–984.

2. Harris JA, Mihalas S, Hirokawa KE, et al. Hierarchical organization of cortical and thalamic connectivity. Nature. 2019/11/01 2019;575(7781):195-202. doi:10.1038/s41586-019-1716-z

3. Murphy KR, Farrell JS, Bendig J, et al. Optimized ultrasound neuromodulation for non-invasive control of behavior and physiology. Neuron. 2024;112(19):3252–3266.e5. doi:10.1016/j.neuron.2024.07.002

4. Shine JM, Lewis LD, Garrett DD, Hwang K. The impact of the human thalamus on brain-wide information processing. Nature reviews Neuroscience. Jul 2023;24(7):416–430. doi:10.1038/s41583-023-00701-0

5. Hallett M. Transcranial magnetic stimulation: a primer. Neuron. 2007;55(2):187–199.

6. Woods AJ, Antal A, Bikson M, et al. A technical guide to tDCS, and related non-invasive brain stimulation tools. Clinical neurophysiology. 2016;127(2):1031–1048.

7. Wickwire EM, Geiger-Brown J, Scharf SM, Drake CL. Shift Work and Shift Work Sleep Disorder: Clinical and Organizational Perspectives. Chest. May 2017;151(5):1156–1172. doi:10.1016/j.chest.2016.12.007

8. Wickwire EM. There is no question about it, sleep disorders increase health care costs. J Clin Sleep Med. Oct 1 2021;17(10):1971–1972. doi:10.5664/jcsm.9606

9. Drake CL, Roehrs T, Richardson G, Walsh JK, Roth T. Shift Work Sleep Disorder: Prevalence and Consequences Beyond that of Symptomatic Day Workers. Sleep. 2004;27(8):1453–1462. doi:10.1093/sleep/27.8.1453

10. Nutt D, Wilson S, Paterson L. Sleep disorders as core symptoms of depression. Dialogues Clin Neurosci. 2008;10(3):329–36. doi:10.31887/DCNS.2008.10.3/dnutt

11. Fang H, Tu S, Sheng J, Shao A. Depression in sleep disturbance: A review on a bidirectional relationship, mechanisms and treatment. Journal of Cellular and Molecular Medicine. 2019;23(4):2324–2332. 10.1111/jcmm.14170

12. Staner L. Sleep and anxiety disorders. Dialogues Clin Neurosci. Sep 2003;5(3):249-58. doi:10.31887/DCNS.2003.5.3/lstaner

13. Peter-Derex L, Yammine P, Bastuji H, Croisile B. Sleep and Alzheimer’s disease. Sleep medicine reviews. 2015;19:29–38.

14. Bubu OM, Brannick M, Mortimer J, et al. Sleep, cognitive impairment, and Alzheimer’s disease: a systematic review and meta-analysis. Sleep. 2017;40(1):zsw032.

15. Fitzgerald T, Vietri J. Residual Effects of Sleep Medications Are Commonly Reported and Associated with Impaired Patient-Reported Outcomes among Insomnia Patients in the United States. Sleep Disord. 2015;2015:607148. doi:10.1155/2015/607148

16. Baek H, Pahk KJ, Kim H. A review of low-intensity focused ultrasound for neuromodulation. Biomedical engineering letters. 2017;7:135–142.

17. Murphy KR, Nandi T, Kop B, et al. A practical guide to transcranial ultrasonic stimulation from the IFCN-endorsed ITRUSST consortium. Clinical Neurophysiology. 2025/03/01/ 2025;171:192-226. 10.1016/j.clinph.2025.01.004

18. Di Ianni T, Morrison KP, Yu B, Murphy KR, de Lecea L, Airan RD. High-throughput ultrasound neuromodulation in awake and freely behaving rats. Brain Stim. 2023/11/01/ 2023;16(6):1743-1752. 10.1016/j.brs.2023.11.014

19. Luck SJ, Kappenman ES, Fuller RL, Robinson B, Summerfelt A, Gold JM. Impaired response selection in schizophrenia: evidence from the P3 wave and the lateralized readiness potential. Psychophysiology. Jul 2009;46(4):776–86. doi:10.1111/j.1469-8986.2009.00817.x

20. Treeby BE, Cox BT. k-Wave: MATLAB toolbox for the simulation and reconstruction of photoacoustic wave fields. J Biomed Opt. Mar-Apr 2010;15(2):021314. doi:10.1117/1.3360308

21. Vyazovskiy VV, Tobler I. Theta activity in the waking EEG is a marker of sleep propensity in the rat. Brain research. 2005;1050(1-2):64–71.

22. Aeschbach D, Matthews JR, Postolache TT, Jackson MA, Giesen HA, Wehr TA. Dynamics of the human EEG during prolonged wakefulness: evidence for frequency-specific circadian and homeostatic influences. Neuroscience letters. 1997;239(2-3):121–124.

23. Siclari F, Tononi G. Local aspects of sleep and wakefulness. Current opinion in neurobiology. 2017;44:222–227.

24. Sheehan DV, Lecrubier Y, Sheehan KH, et al. The Mini-International Neuropsychiatric Interview (M.I.N.I.): the development and validation of a structured diagnostic psychiatric interview for DSM-IV and ICD-10. J Clin Psychiatry. 1998;59 Suppl 20:22-33;quiz 34-57.

25. Weintraub S, Dikmen SS, Heaton RK, et al. Cognition assessment using the NIH Toolbox. Neurology. Mar 12 2013;80(11 Suppl 3):S54-64. doi:10.1212/WNL.0b013e3182872ded

26. Cella D, Yount S, Rothrock N, et al. The Patient-Reported Outcomes Measurement Information System (PROMIS): progress of an NIH Roadmap cooperative group during its first two years. Med Care. May 2007;45(5 Suppl 1):S3-s11. doi:10.1097/01.mlr.0000258615.42478.55

27. Allen Institute for Brain S. Allen Human Brain Atlas. 2013;

28. Barry RJ, Kirkaikul S, Hodder D. EEG alpha activity and the ERP to target stimuli in an auditory oddball paradigm. International Journal of Psychophysiology. 2000/12/01/ 2000;39(1):39-50. 10.1016/S0167-8760(00)00114-8

29. Snipes S, Krugliakova E, Meier E, Huber R. The theta paradox: 4-8 Hz EEG oscillations reflect both sleep pressure and cognitive control. Journal of Neuroscience. 2022;42(45):8569–8586.

30. Hillard B, El-Baz AS, Sears L, Tasman A, Sokhadze EM. Neurofeedback training aimed to improve focused attention and alertness in children with ADHD: a study of relative power of EEG rhythms using custom-made software application. Clin EEG Neurosci. Jul 2013;44(3):193–202. doi:10.1177/1550059412458262

31. Zawiślak-Fornagiel K, Ledwoń D, Bugdol M, et al. The Increase of Theta Power and Decrease of Alpha/Theta Ratio as a Manifestation of Cognitive Impairment in Parkinson’s Disease. J Clin Med. Feb 16 2023;12(4)doi:10.3390/jcm12041569

32. Cavanagh JF, Frank MJ. Frontal theta as a mechanism for cognitive control. Trends Cogn Sci. Aug 2014;18(8):414–21. doi:10.1016/j.tics.2014.04.012

33. Herrmann CS, Knight RT. Mechanisms of human attention: event-related potentials and oscillations. Neuroscience and biobehavioral reviews. Aug 2001;25(6):465–76. doi:10.1016/s0149-7634(01)00027-6

34. Peng W, Hu L, Zhang Z, Hu Y. Causality in the association between P300 and alpha event-related desynchronization. PLoS One. 2012;7(4):e34163. doi:10.1371/journal.pone.0034163

35. Dinges DF, Pack F, Williams K, et al. Cumulative sleepiness, mood disturbance, and psychomotor vigilance performance decrements during a week of sleep restricted to 4-5 hours per night. Sleep. Apr 1997;20(4):267–77.

36. Jurcak V, Tsuzuki D, Dan I. 10/20, 10/10, and 10/5 systems revisited: Their validity as relative head-surface-based positioning systems. NeuroImage. 2007/02/15/ 2007;34(4):1600- 1611. 10.1016/j.neuroimage.2006.09.024

37. Caulfield KA, Fleischmann HH, Cox CE, Wolf JP, George MS, McTeague LM. Neuronavigation maximizes accuracy and precision in TMS positioning: Evidence from 11,230 distance, angle, and electric field modeling measurements. Brain Stimul. Aug 27 2022;15(5):1192–1205. doi:10.1016/j.brs.2022.08.013

38. Scrivener CL, Reader AT. Variability of EEG electrode positions and their underlying brain regions: visualizing gel artifacts from a simultaneous EEG-fMRI dataset. Brain and Behavior. 2022/02/01 2022;12(2):e2476. 10.1002/brb3.2476

39. Oostenveld R, Fries P, Maris E, Schoffelen J-M. FieldTrip: Open Source Software for Advanced Analysis of MEG, EEG, and Invasive Electrophysiological Data. Computational Intelligence and Neuroscience. 2011/01/01 2011;2011(1):156869. 10.1155/2011/156869

40. Mitra PP, Pesaran B. Analysis of dynamic brain imaging data. Biophys J. Feb 1999;76(2):691–708. doi:10.1016/s0006-3495(99)77236-x

