## Supplemental Sections 1-4 for "Focused ultrasound to the bilateral thalamus causally modulates human cognitive attention in a frequency- and intensity-dependent manner"

### Supplemental Section 1: Tolerability Findings

*Side Effects: High Tolerability Across All Stimulation Conditions*

In addition to primary behavioral and EEG measures of interest, we collected tolerability data on each stimulation day at pre-, during-, and post-LIFU timepoints (i.e., 40 minutes after stimulation following all behavioral and EEG tasks). We asked participants to rate headache, pain, skin/scalp irritation, facial twitching, fatigue, and audible clicking noise on VAS scales of 0 (none) to 10 (extreme). Results are summarized in **Supplemental Figure 1**. In most stimulation conditions and ratings, scores were near 0 out of 10 and did not significantly differ. In one exception, for the 1Hz condition participants reported minor skin/scalp irritation, $F(1.02, 15.41) = 6.12$, $p = 0.025$. However, this did not survive post-hoc Tukey corrections for multiple comparisons. In addition, 1Hz LIFU also elicited reduced ratings of fatigue, $F(1.22, 18.23) = 6.85$, $p = 0.013$, with the significant difference being driven by pre-LIFU ratings of higher fatigue ($1.38 \pm 0.38$) as compared to during-LIFU ($0.19 \pm 0.14$). Given that the 1Hz condition also elicited feelings of skin/scalp irritation whereas other conditions caused less of this sensation, perhaps the more pronounced stimulation sensation subjectively caused participants to briefly increase alertness. Lastly, participants reported hearing an audible clicking noise from the US transducers during-LIFU for all stimulation conditions, but not at pre- or post-LIFU timepoints; 1Hz: $F(1.03, 15.37) = 21.66$, $p = 0.0003$, 5Hz: $F(1, 15) = 26.78$, $p = 0.0001$, 25Hz: $F(1, 15) = 21.43$, $p = 0.0003$, and sham: $F(1.02, 15.25) = 34.68$, $p < 0.0001$. Therefore, as all conditions including sham elicited similar during-LIFU audible clicking, such that the unfocused diffuse sham appears to have appropriately replicated the noise that active LIFU stimulation makes and thus mimics the active stimulation conditions.

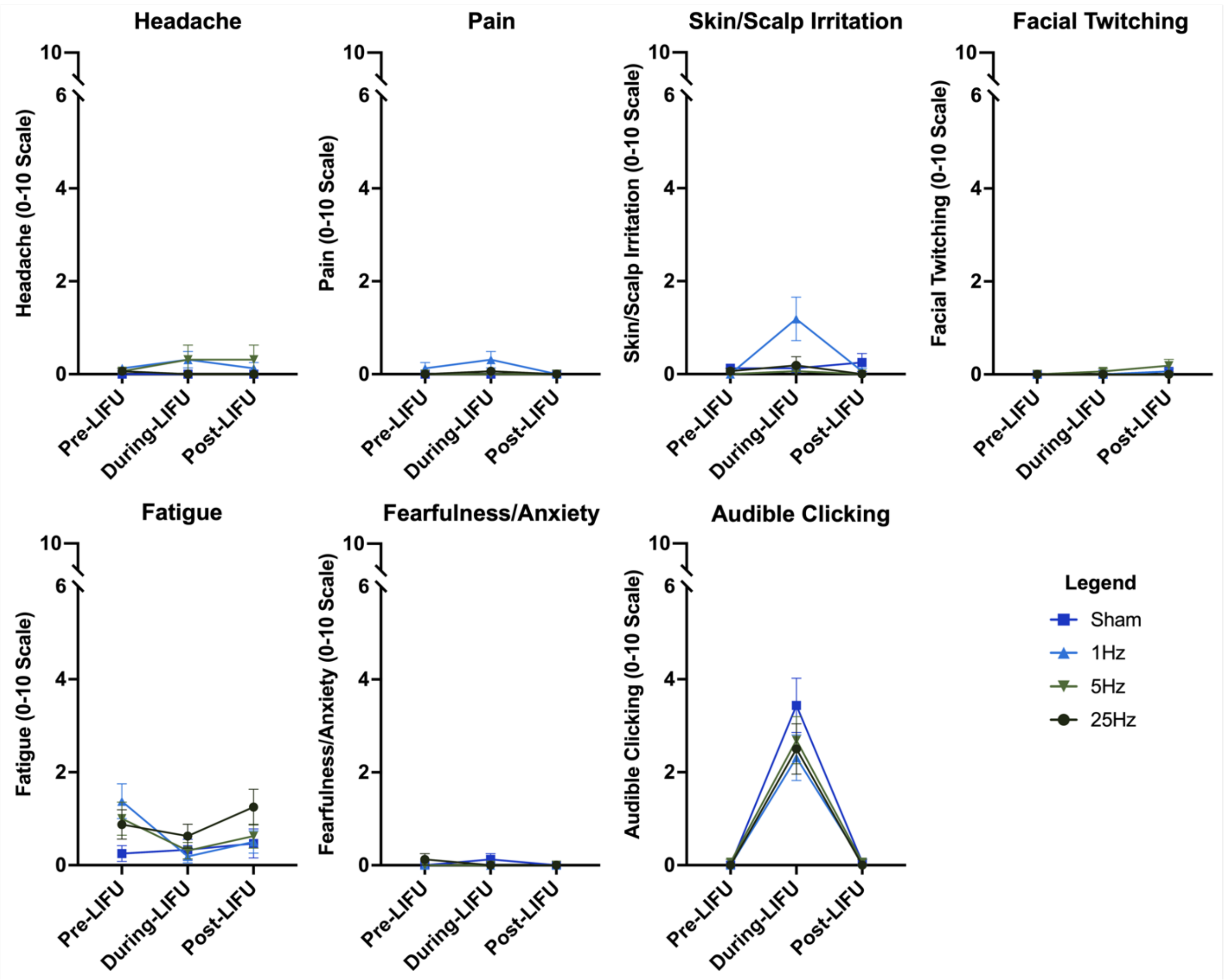


**Supplemental Figure 1: LIFU Tolerability Profile.** LIFU was well-tolerated across 1Hz, 5Hz, 25Hz, and sham stimulation conditions, with minimal ratings of skin/scalp irritation and audible clicking during LIFU that subsided at a post-LIFU timepoint 40 minutes after stimulation. Error bars are SEM.

**Supplemental Section 2: Psychomotor Vigilance Task (PVT) Results**

In addition to the Oddball Task assay of directed attention, participants also completed a Psychomotor Vigilance Task (PVT) that tested their ability to sustain attentional vigilance throughout a 10-minute period. In the PVT, participants click a computer mouse button as quickly as possible after onset of millisecond counters that start at 0ms and are presented every 1-10 seconds. As in the Oddball task, RT is examined in the PVT as a primary indicator of attention engagement; and consistent with task conventions[20] we only included trials where participants responded correctly between 150-500ms post-stimulus onset. Supporting this focus, in this study participants had few lapses (i.e., RTs > 500ms), averaging 0.7 ± 0.28 per administration, and also few false starts (i.e., responses in the inter-stimulus interval), averaging 0.9 ± 0.26.

For PVT, we did not find any pre- to post-LIFU changes in RT for 1Hz, $t(15) = 0.5$, $p = 0.47$, $d = 0.15$, 5Hz, $t(15) = 1.0$, $p = 0.65$, $d = 0.29$, 25Hz, $t(15) = 1.4$, $p = 0.35$, $d = 0.40$, or sham, $t(15) = 0.3$, $p = 0.78$, $d = 0.11$, conditions. This was also true for an evoked theta burst, with no significant LIFU-mediated changes after 1Hz, $t(14) = 0.2$, $p = 0.97$, $d = 0.01$, 5Hz, $t(14) = -0.2$, $p = 0.93$, $d = -0.03$, 25Hz, $t(14) = 1.1$, $p = 0.25$, $d = 0.31$, or sham stimulation, $t(14) = 1.1$, $p = 0.24$, $d = 0.32$. Regarding alpha blocking there were significant pre-to-post increases in the 25Hz, $t(14) = 2.6$, $p = 0.02$, $d = 0.66$, and sham, $t(14) = 3.6$, $p = 0.003$, $d = 0.92$, conditions and a marginal trend in the 5Hz condition, $t(14) = 1.8$, $p = 0.09$, $d = 0.47$. In contrast, an increase in alpha blocking was in the same direction but not significant in the 1Hz condition, $t(14) = 1.5$, $p = 0.15$, $d = 0.39$.

Lastly, we did not find any significant changes for pre-trial frontal theta activity in the 1Hz, $t(14) = 0.1$, $p = 0.99$, $d = 0.03$, 5Hz, $t(14) = 0.5$, $p = 0.66$, $d = 0.12$, 25Hz, $t(14)= -1.4$, $p = 0.17$, $d = -0.37$, or sham, $t(14) = 1.9$, $p = 0.08$, $d = 0.48$, conditions. Conversely, for pre-trial parietal alpha there were significant or trending increases in the 1Hz, $t(14) = 2.6$, $p = 0.02$, $d = 0.67$, 5Hz, $t(14) = 1.9$, $p = 0.07$, $d = 0.50$, and sham conditions, $t(14) = 2.6$, $p = 0.02$, $d = 0.68$, but a non-significant change after 25Hz stimulation, $t(14) = 1.1$, $p = 0.30$, $d = 0.23$.

**Supplemental Section 3: Neurocognitive and Global Health Profile**

We administered the Patient-Reported Outcomes Measurement Information System (PROMIS) and NIH Cognition Toolbox on Visits 1 and 6 to assess general impacts of repeated LIFU on neurocognitive function and global health metrics. We found that participants had below average anxiety scores at Visit 1 (45.63 ± 7.44) which did not change after 4 sessions of LIFU (45.86 ± 7.53), $t(15) = 0.16$, $p = 0.88$, $d = 0.04$; and, depression scores were also below average at Visit 1 (42.87 ± 7.27) and again at Visit 6 (41.98 ± 6.64), $t(15) = 0.59$, $p = 0.56$, $d = 0.15$. Further, there were also no changes in PROMIS-assessed cognition between Visit 1 (53.49 ± 6.67) and Visit 6 (52.3 ± 5.01), $t(15) = 1.04$, $p = 0.32$, $d = 0.26$, nor were there changes in fatigue (47.17 ± 8.02 to 45.03 ± 6.13), $t(15) = 1.17$, $p = 0.26$, $d = 0.29$. Finally, for sleep, there was no significant change in PROMIS T scores, $t(14) = 1.5$, $p = 0.16$, $d = 0.38$, even while it should be noted that participants were above the US average in sleep disturbance at baseline (53.4 ± 7.22) and then below the average at post-LIFU (45.8 ± 7.33).

For the NIH Cognition Toolbox, only 13 of 16 participants had complete datasets. For crystallized cognition, there were no significant improvements from Visit 1 (58.54 ± 8.75) to Visit 6 (61.31 ± 9.96), $t(12) = 1.61$, $p = 0.14$, $d = 0.46$. Conversely, for fluid cognition there were significant improvements consistent with practice effects from Visit 1 (63.0 ± 8.08) to Visit 6 (69.54 ± 8.25), $t(12) = 3.10$, $p = 0.01$, $d = 0.89$. Collectively, there were slight increases in NIH Cognition Toolbox scores between Visits 1 and 6 and participants certainly did not worsen as a result of repeated LIFU stimulation.

**Supplemental Section 4: Oddball Non-Target Trial Reaction Times**

To test for LIFU-elicited changes in directed attention, we utilized the Visual Oddball Task at pre- and post-LIFU timepoints and examined reaction time (RT) changes from each LIFU frequency on target trials. For reaction time, we only included trials with correct responses, and we removed outlier RTs in accordance with standard procedures. Participants were generally accurate, with an 88.3% ± 1.6% rate for target trials and 98.3% ± 0.5% rate for non-target trials.

This supplemental section reports the results of the non-target "distractor" trials, which are also summarized in **Supplemental Figure 2**. We found significant improvements in reaction time from pre- to post-LIFU in multiple conditions that were consistent with practice effects:

Sham: $t(15) = 4.42$, $p = 0.0005$, $d = 1.10$. Reduction of 14.82 ± 3.36ms.
1Hz: $t(15) = 2.19$, $p = 0.045$, $d = 0.55$. Reduction of 11.25 ± 5.14ms.
25Hz: $t(15) = 6.14$, $p < 0.0001$, $d = 1.54$. Reduction of 23.26 ± 3.79ms.

However, there were no significant decreases in reaction time for the 5Hz condition: $t(15) = 1.01$, $p = 0.33$, $d = 0.25$. Reduction of 5.37 ± 5.34ms.

While we were primarily interested in the changes in target reaction times as a measure of cognitive arousal/attention, these distractor data complement the main text in several ways: 1) In contrast to the target data, in which only 25Hz LIFU elicited significant behavioral gains, we observed broader improvements in reaction time across several LIFU frequencies including sham, which appears consistent with practice effects. 2) 5Hz had the lowest behavioral gains and performance improved less than unfocused diffuse sham, potentially pointing to a slight decrease in performance from 5Hz LIFU. 3) 25Hz LIFU elicited the largest improvements in reaction time, possibly corroborating the idea that this higher stimulation frequency leads to the greatest attentional gains.

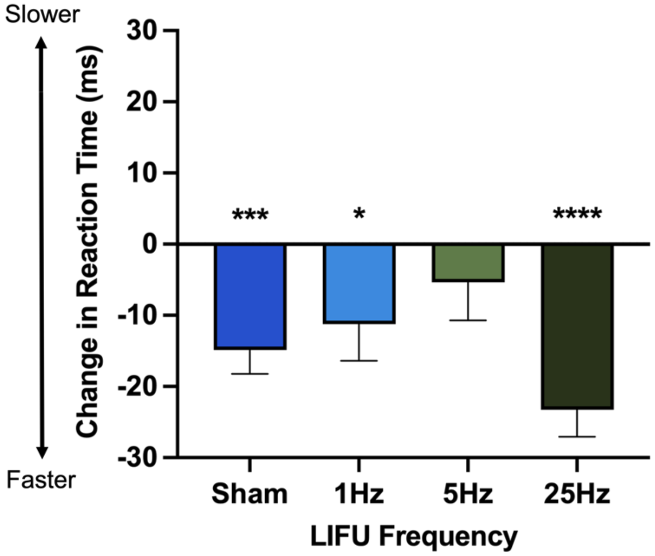


**Supplemental Figure 2: Visual Oddball Non-Target Trial Reaction Times.** There were significant improvements in reaction time across the sham, 1Hz, and 25Hz conditions. *$p < 0.05$, ***$p < 0.001$, ****$p < 0.0001$.
